# What is known about modified insects for disease prevention?: a systematic review

**DOI:** 10.1101/494849

**Authors:** Nazareth Teresa, Craveiro Isabel, Moutinho Alanny, Gonçalves Cátia, Gonçalves Luzia, Teodósio Rosa, Carla A Sousa

## Abstract

The modification and release of insects to suppress or replace natural insect vectors constitutes a promising tool for vector control and disease prevention, facing the unprecedented global emergence of vector-borne diseases. Little is known regarding these innovative modification strategies and available evidence is not standardized turning it difficult to reflect on their actual efficacy and eventual effects.

This work conducted a systematic review, gathering and analyzing research articles from PubMed and Biblioteca Virtual em Saúde databases whose results directly report efficacy and effects of the use of modified insects for disease prevention until 2016. Within more than 1500 publications that were screened a total of 349 where analyzed.

A total of 12/3.4% reported field-based evidence, and 41/11.7% covered modification stretagies’ efficacy after insects’ release, their epidemiology impact or its long-term efficacy. Examples of successful results were the replacement of natural field populations by *wolbachia-infected* mosquitoes in 5 weeks, and the elimination of a population in laboratory cages after transgenic mosquitoes release over 10–20 weeks. Variability in the effective results were described (90/25.7%) questioning its reproducibility in different settings. We also found 38/10.9% publications reporting reversal outcomes, such as an increase of vector population after release. Ecological effects such as horizontal transfer events to non-target species (54/15.5%), and *wolbachia*-induced worsening pathogenesis on mammal filarial diseases (10/2.9%) were also reported.

Present work revealed promising outcomes of both suppressing and replacing approaches. However, it also revealed a need of field-based evidence mainly regarding epidemiologic and long-term impact of insect modification strategies. It pointed out some eventual irreversible and important effects that must not be ignored when considering open-field releases, and that may constitute constraints to generate the missing field evidence. Moreover, the level of variability of existing evidence suggests the need of local/specific evidence in each setting of an eventual release.

**Author summary:** Innovative strategies are needed to arrest the unprecedented increase of vector-borne disease incidence, distribution and severity. Several modification techniques are being tried all over the world. However this is still an emergent topic with scarce available information and of complex understanding.

Present work is the unique structured review regarding the use of modified insects for vector-borne disease prevention, bringing neutral and robust evidence that will contribute with critical insights regarding these approaches.

Here we explored more than 1200 publications and analyzed 349 publications on this subject, describing the actual efficacy and reported effects of several modification strategies. More than 30 categories were reported such as, the type of modification, the year of the publication, the species were results were tested, the type of study, and also type of Efficacy Outcome (from modification to long-term) and/or the type of Effects Outcome (from physiologic to ecologic effects). Analysis revealed promising outcomes regarding vector-control and disease prevention. However insects’ modification strategies still lack field-based evidence mainly regarding epidemiological and long-term efficacy. Eventual reversal outcomes on disease transmission, or irreversible biological effects (including horizontal transfer to non-target species or worsening pathogenesis in particular diseases in mammals), were also described. These effects need to be explored, dispelled or resolved before field trials occur in human residential areas. Some of these questions could only have a robust answer if these strategies would be implemented, needing to take the risk to observe reversal outcomes and/or irreversible effects. These findings reflect the big dilemma that is under the use of modified insects to prevent vector-borne diseases. Findings could support health authorities in decision-making and regulatory committees during advisory processes, by evaluating the pros/cons of each modifying technique for a particular setting. Moreover they could also summarize what is crucial to inform to communities if planning open field releases in residential areas.

## INTRODUCTION

Vector-borne diseases have a wide impact on human health being an mandatory topic on global health agendas (1)(2). Even with the significant reduction of the global burden of malaria since the beginning of the century, in 2016, this infectious disease was still responsible for 445 000 deaths (3). Due to human population growth, globalization, and climate change, arboviral diseases outbreaks have been increasing in frequency, expansion, diversity and severity (4). Only dengue’s incidence grew more than 30-fold in the last 50 years (5). Although arboviruses dispersal is partially conditioned by the environmental constraints that limit the distribution of its main vectors, outbreaks of diseases such as, yellow fever, chikungunya and Zika have been reported all over the world (6-10). The severity of Zika fetal malformations during 2015/2016 epidemics turn it a public health emergency of international concern according to World Health Organization (11). The lack of effective approved vaccines for some of these infections and the increase of insecticide resistance in its most competent vectors, impose an urgent need for innovative effective strategies to minimize these diseases (12)(13).

The release of modified insects is considered a promising approach for prevention and control of vector-borne diseases. Innumerous techniques and insects’ modification strategies had been laboratorial tested, all of them fitting one of the two broad approaches: (i) modification and release of sterile insects aiming the reduction/eradication of natural vector populations (suppression approach / vector control approach) or (ii) modification and release of insects refractory to pathogen transmission aiming the replacement of natural vector populations (replacement approach / transmission prevention approach). Open releases of modified insects have been occurring all over the world in an attempt to cope to the unprecedented vector-borne diseases burden (14-21). However, none of these modifying technologies has yet been approved by the WHO’s Vector Control Advisory Group(22).

Few studies reported the effectiveness of insects modification strategies, and even less their eventual effects exploring them only barely and theoretically(23-25). Important reviews on this topic were recently published, but corresponding to the perspective of the author regarding the subject or a summary of the authors’ selection of publications (23)(26-29). This work presents a unique structured review on the use of modified insects to control and prevent vector-borne diseases, gathering, exploring, and classifying evidence available up to 2016 regarding efficacy and effects of the use of modified insects for disease prevention.

## METHODS

Present work is enrolled in a bigger project whose aim is the description of the strengths/weaknesses/opportunities/threats of modified insects for disease prevention. During analysis and reviewers’ consensus, it was realized that all evidence found constituting strengths/weaknesses/opportunities/threats of the insects’ modification for disease prevention, were fitting in two main themes: the efficacy and the effects of the insects’ modifications. Hence, results were extracted and are presented according to this classification.

### Search strategy

To identify relevant documents focusing on strengths, weaknesses, opportunities and threats of modifying insects to prevent diseases, two electronic databases (PubMed and Biblioteca Virtual em Saúde, BVS) were searched using combinations of MeSH terms and free text words such as: “organisms, genetically modified” (MeSH), “wolbachia”, “lethal”, “sterile insect”, “vector-borne”, “replacement” and “suppression”. To help increase sensitivity and specificity, combinations of different search strings were used for each electronic database. Results from all searches were downloaded into Mendeley program (Elsevier); duplicates were withdrawn automatically using Mendeley and verified manually, followed by the inclusion process implementation.

### Study selection

Publications were included in the study when all of the following inclusion criteria were met:

1. Research articles *i.e.* publications structured as Introduction, Material and Methods, and Results/Discussion, or similar.
2. Available as Free Full-Text at NOVA Discovery platform
3. Written in English, French, Portuguese or Spanish.
4. Published until the date of the search (1^st^ March 2016)
5. Publications covering modified insects or the modifications itself. It was considered ‘modification’ any process, species or condition, described in the literature as able to be used to modify insects (rather genetic or other type of modification), even if not explicit in the collected paper.
6. Publications whose results explicitly report strengths, weaknesses, opportunities and threats of the modifications concerned with regard to health impact and/or biological impact
7. Publications whose studies were performed in Insect or Mammal species, rather *in vitro, in vivo, ex vivo* or in archetypal modeled species.

A two-stage inclusion process was applied. All references were initially screened by title and abstract and included in the study if they met the selection criteria. In the second stage, the full text reading of each publication was undertaken. To establish consensus in criteria application, part of the publications (5% of the 1^st^ screening, and 50% of the 2^nd^ screening) were screened by two reviewers (inter-reviewer check). Disagreements were resolved by discussion. In the end of the 2^nd^ screening and after criteria had been discussed, the full-text screening was repeated by one reviewer to ensure homogeneity of the criteria during the process (temporal check). All documents considered relevant went to the next phase of extracting data and analysis.

### Data synthesis and analysis

Data was extracted from the included publications into a digital data-extraction form. Two investigators performed data extraction and analysis of 50 % of the included publications (inter-reviewer check). All extracted data was structured into two major themes, efficacy and effects of the modification strategies, and each of them divided into several topics and sub-topics (see more detailed information in Results section). These hierarchical categories were defined by the two reviewers through a consensual process. Disagreements were resolved by discussion. When consensus was attained, categorization and analysis of all included articles were re-checked in order to ensure a homogeneous analysis (temporal check). According to the evidence reported, publications were classified into as many categories as possible, in order to reduce the likelihood of missing key points in the data. It was also extracted information regarding the year, type of study (laboratory, semi-field, field and computational modelling), species involved in the experiments, and modification strategy (*Wolbachia*, anti-pathogen Transgenesis, lethal Transgenesis, etc.). As to *Wolbachia*-based studies, publications were classified according to the endosymbiont origin: natural occurrence, artificially introduced or removed from natural or artificially infected hosts. The classification regarding the type of study was used as a *proxy* of the publication robustness, considering semi-field and field studies the most robust ones, and computational modelling and laboratorial the less robust. Apart from qualitative analysis, descriptive statistics analysis was performed. The softwares Excel (Microsoft Office, Windows 10) and NVivo 10 (QSR International Pty Ldt, Doncaster, Victoria, Australia) were used during the analysis. Results are presented by theme, modification strategy and species, publications are also referred according to their chronological order on the manuscript’ sections, tables and supplementary information. This literature review followed the proceeding of a PRISMA methodology (S1 Checklist).

## RESULTS

Databases searches resulted in a total of 1567 publications (Fig 1). Following the removal of duplicates, 1205 references were selected. After the two-stage selection process 377 articles were included in the study, and 349 publications were analyzed. References from analyzed publications were ordered from 1 up to 349 and cited in *italic* for differentiation from manuscript’s references (see full list of analyzed publications and summary of analysis in S1 Appendix).

**Fig 1.**
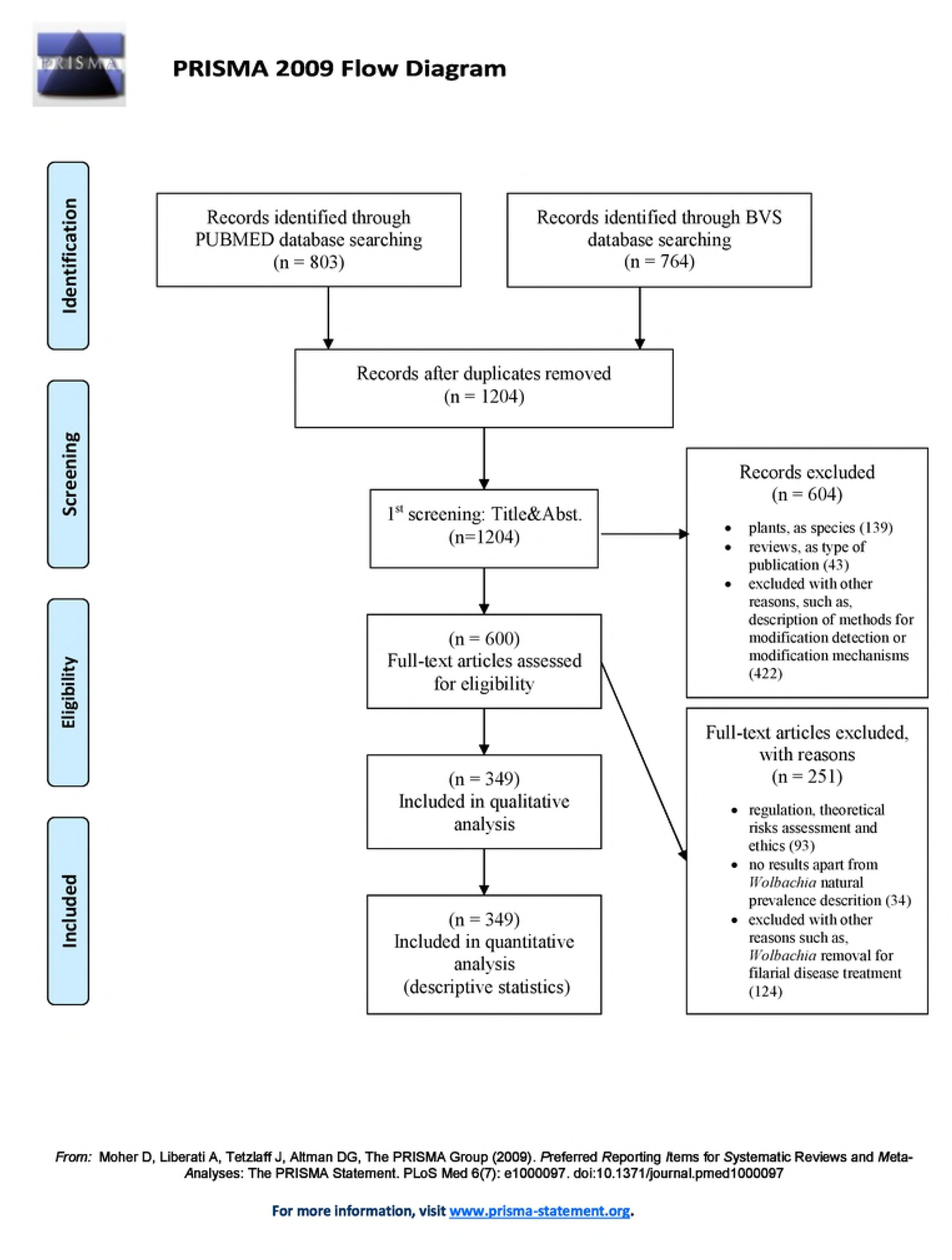
PRISMA Flowchart reporting the number of publications in each stage of the review

From the 349 publications selected for analysis, 340/97.1% were published after the year of 2000. The majority constituted laboratory studies, *i.e*. performed in a controlled experimental environment (310/88.6%) and referred to *Wolbachia* or other symbiont-based modification strategy (307/88.0%). Out of those 307, 2/0.7% publications referred to other symbiont (*Rickettsia* and *Sodalis*) *(1),(2)*. Several organisms’ species integrated in the experiments of the analyzed publications: five genera of insects vectors (*Aedes, Anopheles, Culex, Mansonia, Phlebotomos* and *Glossina*) 54 genera of non-vector insects, and four mammals genera (see all data regarding quantitative analysis in S1 Figure).

In what concerns the content of the publications, two major themes emerged from the analyzed research articles: (i) efficacy of the modification strategies, and; (ii) effects induced by the modifications. Both themes (efficacy and effects) were divided into several topics (see Fig 2).

**Fig 2.**
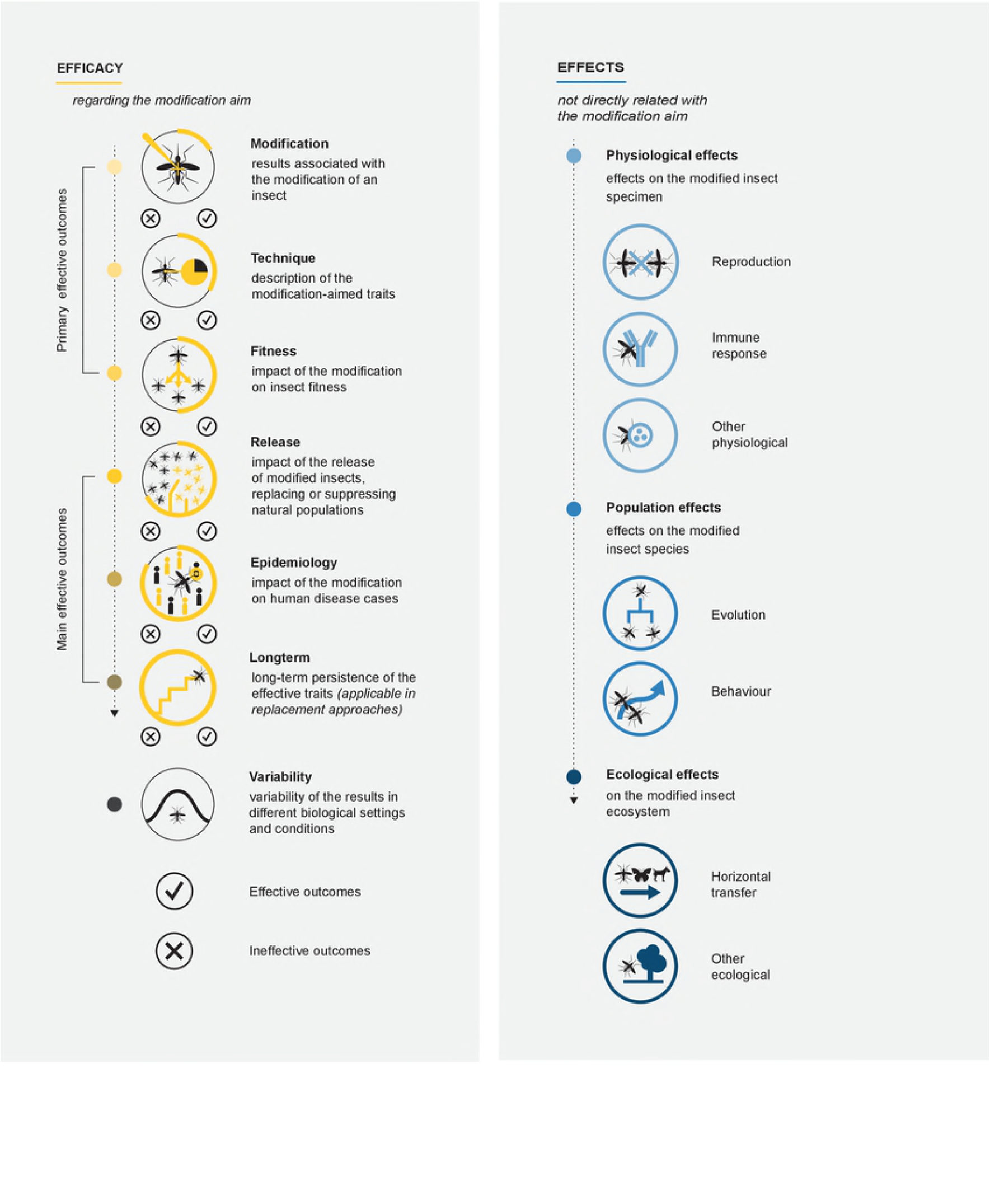
Schematic representation of the themes, topics and type of outcomes described in the Results’ section

Efficacy outcomes were also divided into effective outcomes - reporting the success of the modification strategy - and ineffective outcomes - reporting the failure of the modification strategy. Ineffective outcomes includes outcomes that: (i) achieved no results, (ii) described indirect results that call into question the efficacy of the modification strategy and/or (iii) reported reversal results, i.e. that lead to the reverse of the aim of the modification strategy. There were more publications contributing to efficacy (237/67.7%) than publications with results regarding effects (156/44.6%) (Fig 3). Regarding themes and topics’ analysis, each publication may have contributed to more than one category.

**Fig 3.**
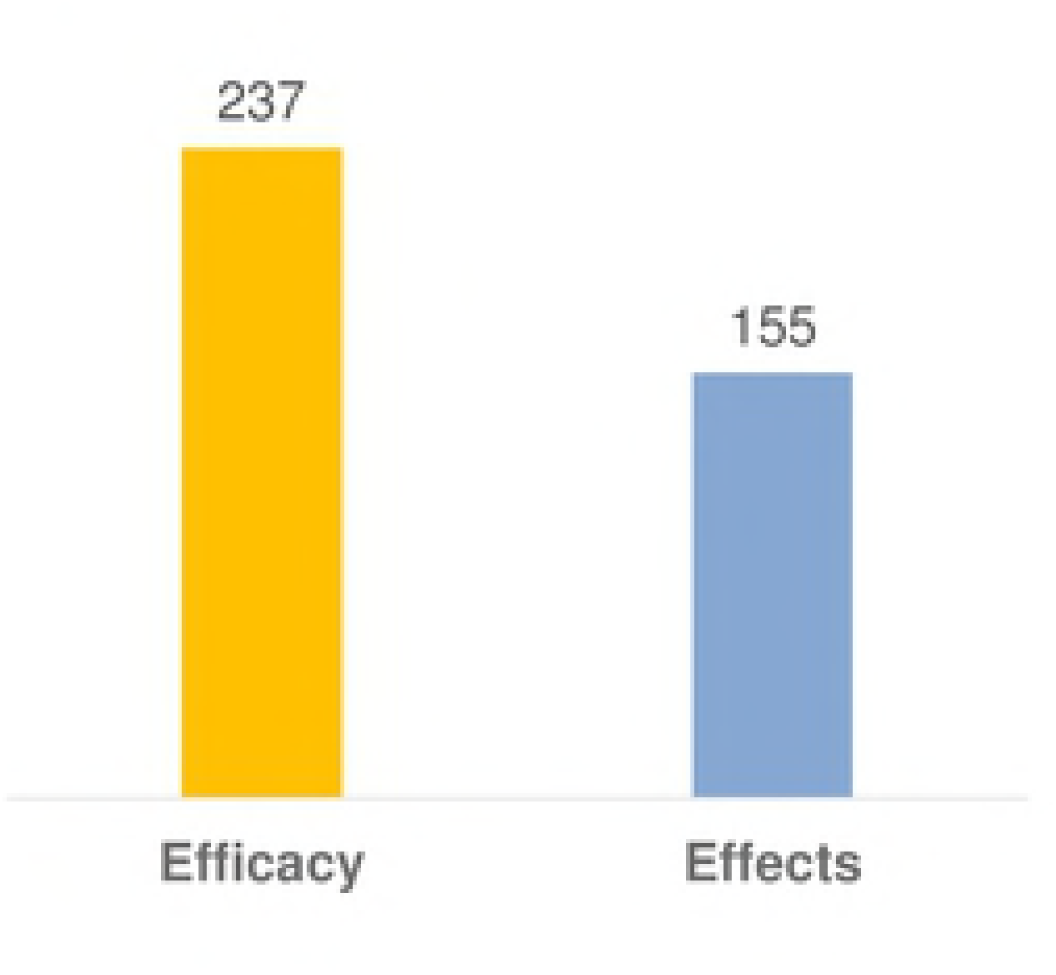
Distribution of the publications whose results contributed to each of the major themes (n=349)

In what concerns publications covering efficacy outcomes, there were more publications reporting ineffective outcomes (164/69.2%) than publications reporting effective ones (150/63.3%). Out of the latter, only 41/27.3% constitute main effective outcomes (regarding its release, epidemiologic and long-term efficacy), and out of the former, 38/23.2% constitute reversal outcomes (Fig 4).

**Fig 4.**
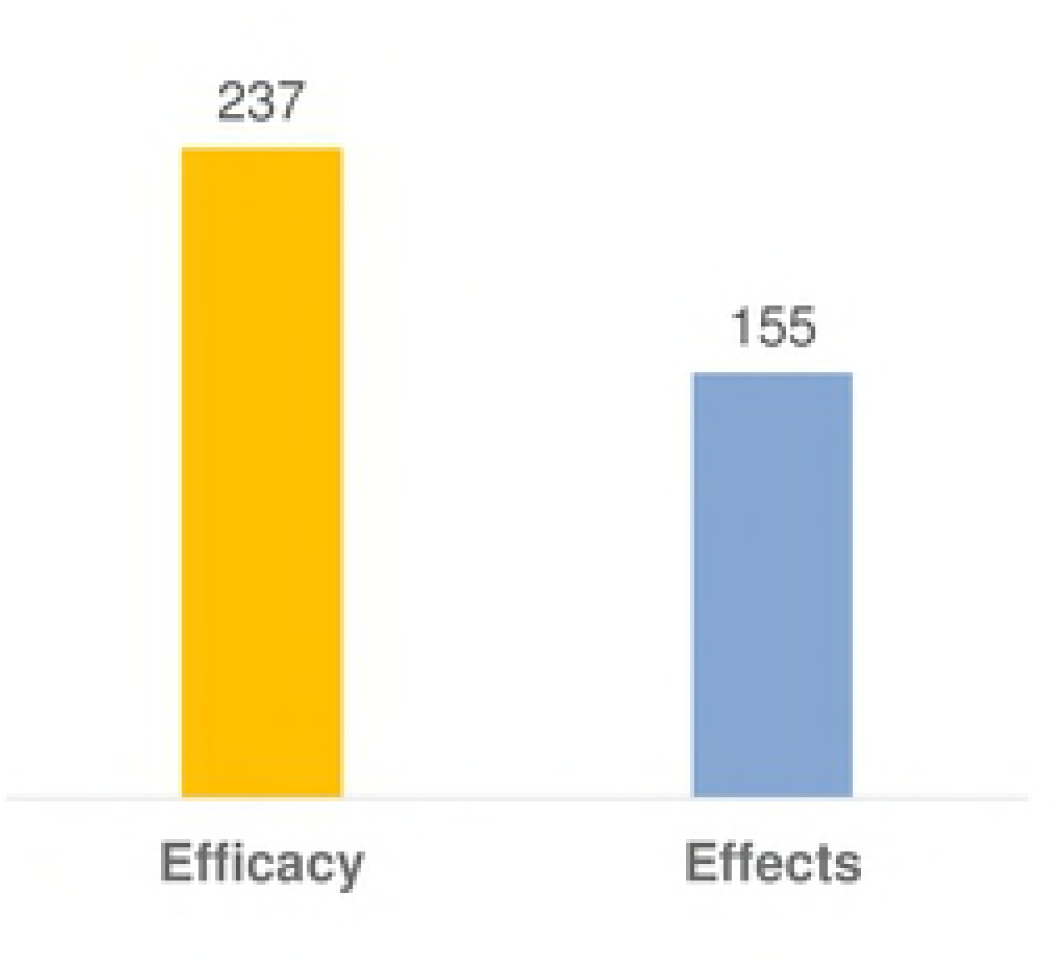
Distribution of the publications whose results contributed to each type of outcomes in Efficacy (n=237)

Next sections summarize the analysis by themes and topics, and by modification strategies as follows: (i) *Wolbachia* and other symbiont-based modification strategies and (ii) Transgenesis and other non-symbiont-based modification strategies. Some modifications are the combination of the two above types of modification strategies. Examples of those are: (i) Paratrangenesis, *i.e.* the introduction of a transgene in a symbiont bacteria infecting the insect (rather than introducing the transgene in the genome of the insect itself) *(2)*; (ii) the simultaneous introduction of a *Wolbachia* and a transgene in the same organism *(3)* (both included in ‘*Wolbachi*a and other symbiont-based’ section); and (iii) Transgenesis using *Wolbachia* as gene drive *(4-9)* (included in ‘Transgenesis and other’ section).

## EFFICACY

### *Wolbachia* and other symbiont-based modification strategies (effective and ineffective outcomes)

A total of 195/82.3% research articles, out of the 237 with efficacy outcomes, presented results regarding the efficacy of *Wolbachia* and other symbiont-based insect modifications (111/56.9% reported effective outcomes, 145/74.4% reported ineffective outcomes, nTotal=195). One publication reported the efficacy of other symbiont-based insect modification strategy (a *Sodalis* modified by paratrangenesis) *(2)*, and two publications referred to computational studies using archetypal modeled endosymbionts *(10),(11)*. Efficacy results covered all the topics (Modification, Technique, Fitness, Release, Epidemiology, Long-term and Variability) as presented in Fig 5 and described in the following paragraphs.

**Fig 5.**
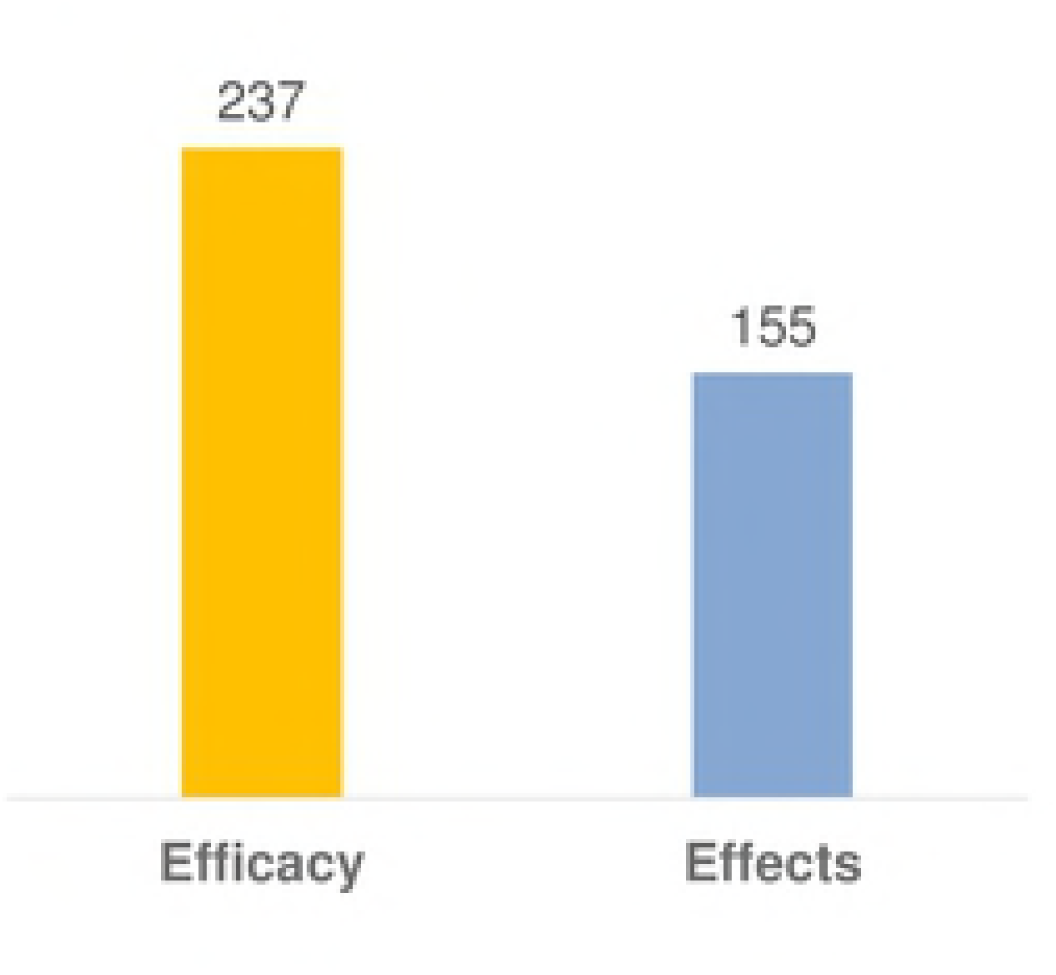
Distribution of the publications whose results contribute to each efficacy topic, distinguishing effective/ineffective outcomes (n=195)

#### • Modification

According to analyzed publications, different *Wolbachia* strains were microinjected into several insect species, leading to a successful germline infection of the insect (stable transinfection). Stable transinfections were reported in the following insect species: *Aedes aegypti (12-31*), *Aedes albopictus (32-36*), *Aedes polyniensis (37),(38), Anopheles gambiae (39*), *Anopheles stephensi (40),(39),(41),* and *Culex tarsalis (42)* (vector mosquitoes), and *Bemisia tabaci* (whitefly) *(43), Ceratitis capitate* (medfly*) (44), Drosophila melanogaster* (fruit fly) *(45),(46), Drosophila simulans* (fruit fly) *(47-50),(45),(51),(46),(52), Ephestia kuehniella* (butterfly) *(53),(54), Laodelphax striatellus* (planthopper) (*55)* (non-vector insects). Apart from transinfections, analyzed articles also described other types of *Wolbachia* successful introduction such as: transient somatic infection (*56-58)*, infections in cell lines *(59-65)*, ex *vivo* organ culture *(66)*, outcrossing *(67),(68)* and introgression *(69-71),(45),(72),(19),(67),(73-77)*. One article reported the loss of *wMe*l on an *Aedes albopictus* cell line 12 passages post infection *(61)*.

#### • Technique

In what concerns the traits that ensure the *Wolbachia* effectiveness, several publications reported high cytoplasmic incompatibility (CI), *i.e.* the survival of offspring from infected females (>90%) and/or perfect (100%) maternal transmission, *i.e.* the transmission of *Wolbachia* to the insect host’s offspring, on vectors species mainly in *Aedine* species *(12),(17),(19),(68),(27),(30),(74),(78-80),(75),(37),* but also in *Anopheles stephensi (40),(39), and Culex pipiens quinquefasciatus (81),(82). Wolbachia*-induced pathogen protection was first reported in *Drosophiline* species, protecting from infection of several RNA virus *(83)*. Later, pathogen protection induced by artificial introduction of *Wolbachia* was also demonstrated in several vector species: not only in those naturally not infected by *Wolbachia*, such as, *Aedes aegypti (14),(16),(19),(20),(28),* and *Anopheles stephensi (40),(84)*; but also in those that naturally host it, such as, *Aedes albopictus (35),(85),(86), Aedes polyniensis (37),(38), Anopheles gambiae (57),(58),* and *Culex quinquefasciatus (87).* In *Aedine* vector species, transinfected *Wolbachia* blocked the development of several human pathogens: yellow fever virus (YF) *(20)*, chikungunya virus (CHIKV) *(16),(20),(88),(86)* three serotypes of dengue virus (DENV1, DENV2 and DENV3) *(16),(19),(28),(64),(35),(85),(38),* and the filarial nematode *Brugia Pahangi (37*). *Wolbachia*-infected *Aedes aegypti* was also protected against *plasmodium gallinaceum (16)*. In *Anopheline* species, transinfected *Wolbachia* induced protection against *Plasmodium falciparum*, the most virulent plasmodium species *(57),(40)*, modestly suppressed *Plasmodium berghei* oocyst levels *(58)*, and at some temperatures also protected from *Plasmodium yoelii* infection *(84)*. In *Culex pipiens quinquefasciatus,* transinfected *Wolbachia* diminished the *West Nile virus* (WNV) transmission (*87*). Six publications also reported that transinfected *Wolbachia* could enhance infection of some pathogens in vector species such as, WNV in *Culex tarsalis (42) and in vitro (59), Plasmodium yoelii* at some temperatures in *Anopheles stephensi (84), Plasmodium gallinaceum in Aedes fluvialitilis (80), Plasmodium relictum in Culex quinquefasciatus (89),* and DENV2 in *Aedes aegypti (90*). These constitute reversal outcomes, *i.e.* the reverse to what was intended with an effective *Wolbachia-*based modification.

#### • Fitness

None or low fitness costs, caused by the introduction of *Wolbachia* in insects species, allow the modified insect to be reproductively competitive against their natural counterparts, and therefore to easily invade natural populations after its release. Analyzed publications described no or low fitness costs after *Wolbachia introduction* in several vector insect species: *Aedes aegypti (67),(22),(24),(73),(30), Aedes albopictus (91),(78),(79),(86), Aedes fluviliatilis (80), Aedes polyniensis (75-77)*, and *Culex pipens quinquefasciatus (81),(82).* Some articles reported a fitness cost that act as part of the control strategy, *i.e*. constituting part of the modification efficacy. These were the report of *Wolbachia*-induced life shortening of vectors (that reduce or eliminate the time vectors can transmit the pathogen) *(16-19),(68),(34),(57)* or decreased viability of desiccated eggs (preventing the next generation of mosquitoes from hatching after the dry season) *(18),(68).* Despite referring to a fitness cost, since they constitute *Wolbachia* traits that ensure its effectiveness, are herein considered effective outcomes of the Technique topic. In some cases *Wolbachia* induced fitness costs on mating competitiveness *(41),* fecundity *(22),(27), (80),(34),(41),* fertility *(22)*, larvae competitiveness *(92)*, life span *(34)*, or development time *(68),(25)* of the modified insect. Moreover, some publications reported *Wolbachia*-induced fitness benefits in vector insects *(13),(90),(93),(74),(92),(41),(94),(95),(82).* Once fitness benefits lead to an increase of the insect vectorial capacity and consequently to an increase on disease transmission, they constitute reversal outcomes.

#### • Release

A successful release of modified insects describe either: (i) their effective invasion and establishment in the field (replacing natural population); or (ii) incompatible mattings between modified and natural insects (suppressing the population). The release of the non-vector *Ceratitis capitata* (fruit fly) males, transinfected and inducing complete CI, led to the complete suppression of a laboratory cage population of natural specimens *(44)*. Insects with introduced *Wolbachia* successfully replaced natural specimens in laboratory cages *(12),(44),* in semi-field cages *(19),(40)*, and in the field, *(67),(27).* Similar results were suggested by computational modelling studies *(96),(30).* Other articles reported invasion but only under certain meterological *(68)* or entomological conditions *(26*), or if some technical ordeals could be overcome *(97)*. However, release of *Wolbachia*–insects also led to no/low invasion rates *(98),(99).* Several studies suggested the need to release prohibitively large number of insects *(100-102).* To overcome that, two solutions were reported: releases in a ratio of 95% male mosquitoes (requiring a mass rear capacity) (*11*) or the introduction of insecticide resistance genes along with *Wolbachia* in the host insect, combined with a pre-release intervention to reduce (adult) insect vector numbers *(29), (3)*. The unintended increase of the insect population after the release of the modified insects (reversal outcomes) was estimated by computational modelling studies *(103)*, some of them based on field data of wMelPop-*aegypti (100)* and of superinfected *Aedes albopictus (93)*.

#### • Epidemiology

No publications described the impact of *Wolbachia*-based modified insects on human disease incidence. However, several computational modelling studies estimated a successful epidemiological impact after the release of *Wolbachia*-insects *(96),(11)*, specifically using wMel-*aegypti* combination which seems to eliminate DENV transmission in low or moderate transmission settings *(104),(105)*.

#### • Long-term

Released wMel-*aegypti* populations persisted in near fixation and maintained the *Wolbachia*-induced DENV protection, two years after the release *(27),(28)*. Laboratory and/or computational modelling studies also revealed that *Wolbachia* (natural or introduced) may persist over time in its symbiotic host *(106),(107),(10),(108).* Despite that, the long-term efficacy of *Wolbachia*-based modification strategy was questioned due to the report of: (i) a change in CI rates or in *Wolbachia* density with age, time or over generations *(109-111),(69),(112),(47),(113-115)*; (ii) a change in other effective traits *(45),(60)*; (iii) loss of *Wolbachia* infection *(106),(116-122)*; and (iv) its natural replacement by other *Wolbachia* strain *(123-125)*. Moreover, long-term efficacy was also questioned by the risk of immigration/re-invasion of other insect populations after the release and fixation of the modified insect population *(21),(27)*. How and how much this phenomena will affect efficacy is not known *(104),(21)*.

#### • Variability

Finally, also affecting *Wolbachia* efficacy is its variability. It was reported that a considerable degree of variability may evolve in short evolutionary periods *(126)*. Several articles described *Wolbachia* evolution *(127),(128)*, including its transition from facultative parasite to a nutritional mutualist *(129)* or obligatory symbiont *(130)*. *Wolbachia* density inside an insect-host changed according to a multitude of factors, such as, host genetic background *(112),(131),(61),(132),(133),(62),(134)*, presence of resistance genes *(135)*, host gender *(109),(136),(111),(134),* development stage *(111)*, nutrition *(57),(41),(137),* immunity status *(111)*, presence of pathogens *(23),(83),* or host microbiome *(39),(138-140).* It also varied according to *Wolbachia* strain *(141),(142)*, even when coexisting in the same host *(140)*, insect larvae density *(143)*, and environmental conditions *(84),(144),(145),(143),(111)* (such as humidity and temperature). However, *Wolbachia* density in *Ae. aegypti* did not alter after repeated human blood feeding *(31)*, neither insecticide susceptibility of *Ae. aegypti* changed after *Wolbachia* infection *(36).* Somewhat surprising *Aedes albopictus* cell lines infected with wStr or wAlbB showed resistance to streptomycin *(63) Wolbachia* main effective outcomes in insect vectors are presented in Table 1 and its reversal outcomes in insect vectors is described in Table 2, see the complete data (also (146-153) and (154-201)) in S1 Table.

**Table 1.**
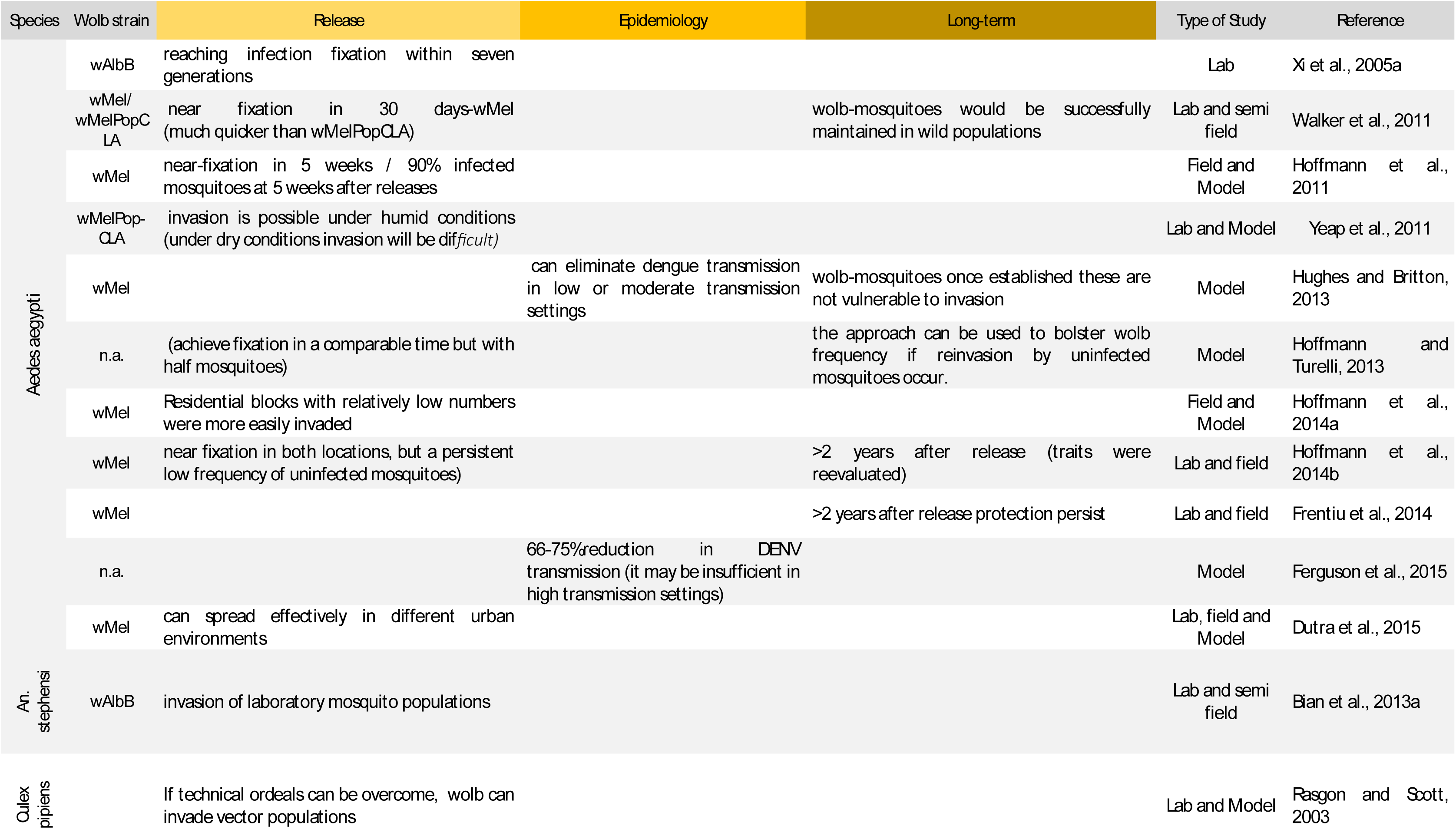
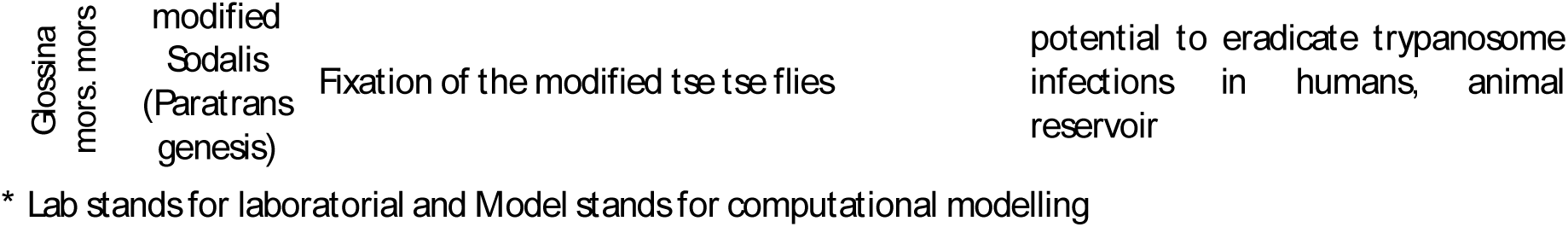
Main effective outcomes of Wolbachia-based and other symbiont-based insect modification (vector insect species). Publications reporting effective outcomes in release, epidemiology and long-term topics (the ones closer to the strategy aim, that is transmission blockage or vector suppressing). Ineffective outcomes of the mentioned publications are also presented (in italic).

**Table 2.**
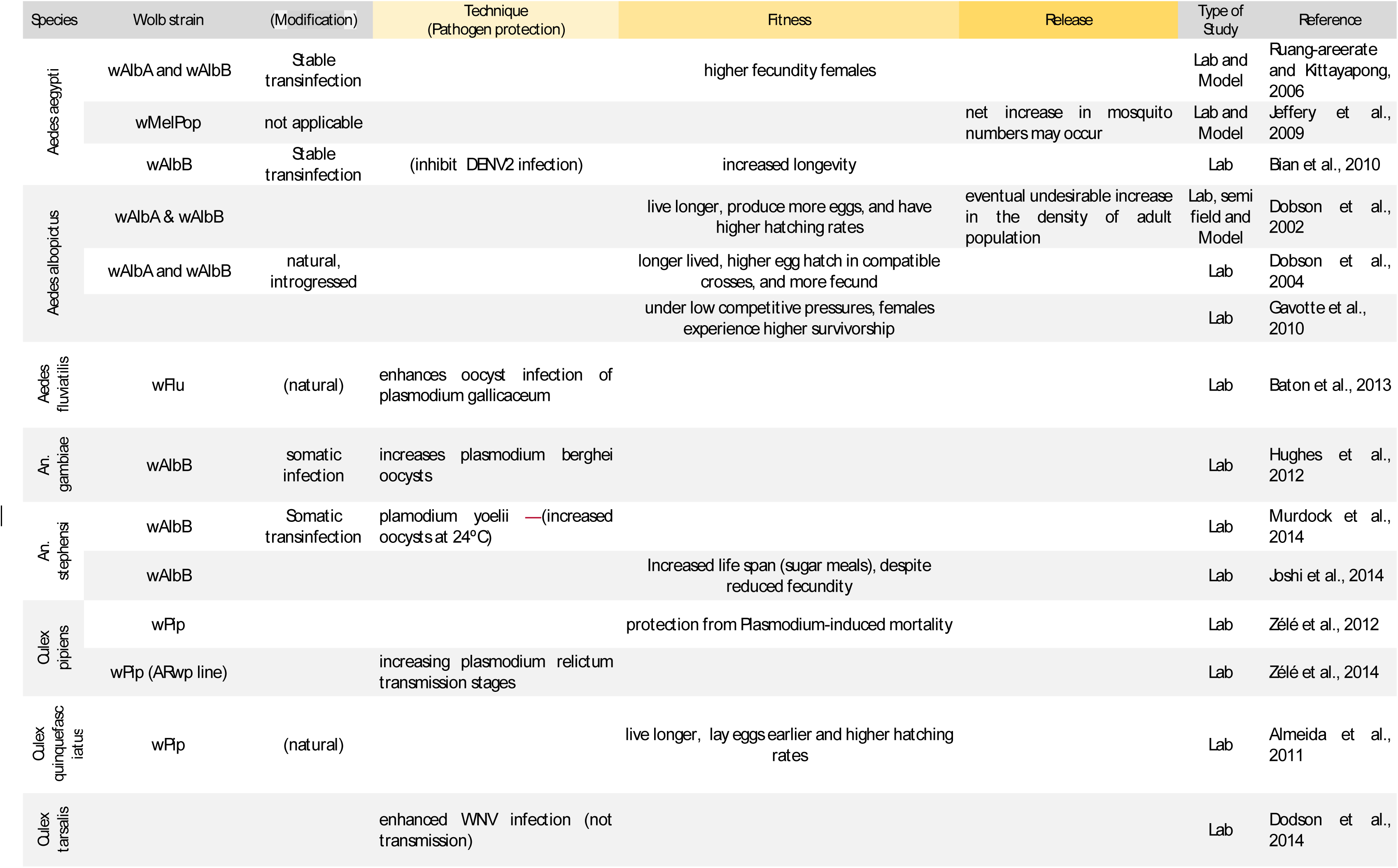
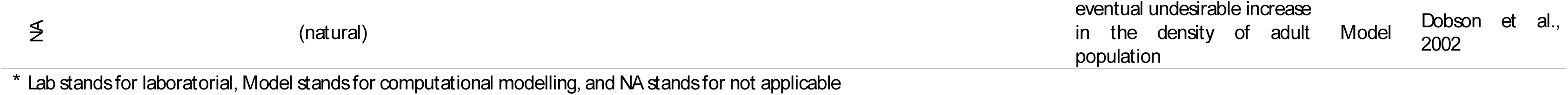
Reversal outcomes of Wolbachia-based and other symbiont-based insect modification (vector insect species cell-lines not included) Publications reporting reversal outcomes (reverse to the strategy aim, that is transmission blockage or vector suppressing). Effective outcomes of the mentioned publications are also presented (in italic).

### Transgenesis and other non-symbiont-based modification strategies (effective and ineffective outcomes)

A total of 37 research articles (15.6% out of the 237 with efficacy outcomes) have results regarding the efficacy of transgenesis or other non-transgenic and non-symbiont-based insect modification (37/100% reported effective outcomes, 15/40.5% reported ineffective outcomes). Two types of trangenesis were found in analyzed publications: (i) anti-pathogen transgenesis, *i.e.* transgenesis with an anti-pathogen effector gene (30 publications*) (202-205),(9),(206),(7),(207), (208),(5),(209-215),(6),(4),(216),(8),(217-225)*; and (ii) results on lethal transgenesis, *i.e.* transgenesis with a lethality inducing gene, covering the release of insects with a dominant (RIDL) (three publications) *(226-228)*, and with a female-killing transgene (three publications) *(229),(224),(230).* Two non-transgenic non-symbiont-based insect modifications were also reported: (i) a radiation-based sterilization insect technique (SIT) (one publication) *(231*), and (ii) a RNAi-mediated sterilization (one publication) *(232*).

Not all modifications covered all efficacy topics (Modification, Technique, Fitness, Release, and Variability). No studies reported results regarding epidemiology efficacy using these types of modifications. Results are described in the following paragraphs and quantified in Fig 6.

**Fig 6.**
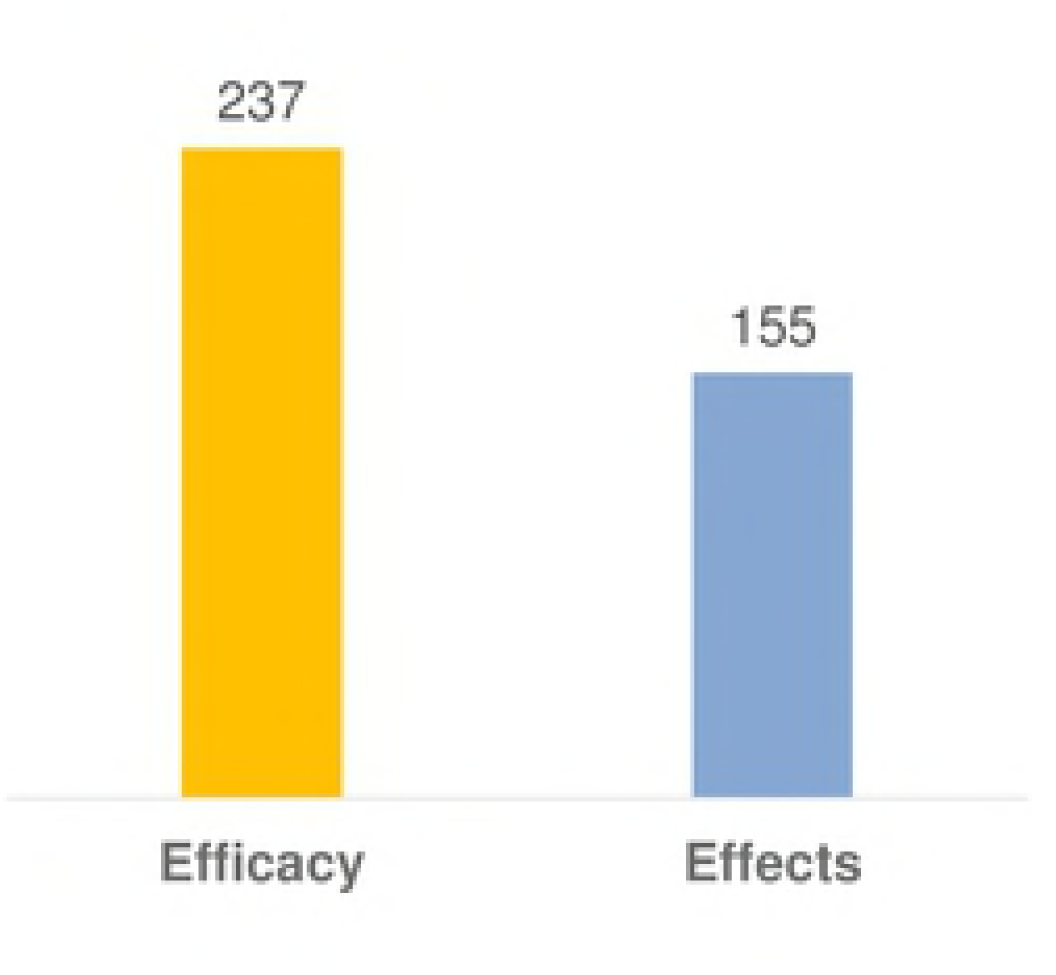
Distribution of the publications whose results contribute to each efficacy topic distinguishing effective/ineffective outcomes (n=37)

#### • Modification (anti-pathogen transgenesis)

Several studies described successful gene vectors mainly transposable elements, such us, *Hermes* and *piggyBac (230),(202),(205),(211),(213),(216),(218),(221),(223),(228),* but also transcription activator-like effector nuclease (TALEN) (225). Moreover, several distinct promotors were described generating successful sex, tissue or stage-specific expression of effector genes *(211),(212),(204),(213),(203),(217),(219),(215).* Successful insertion of an anti-pathogen effector gene was reported in *Aedes aegypti (203),(208),(211),(212),(216),(220),(217),(224), Anopheles gambiae (204),(213),(218),(222),(205),(207),(210),(221), Anopheles stephensi (205),(207),(210),(214),(221), Culex pipiens (7*), G*lossina morsitans morsitans (4)*, and in some non-vector insects *(202),(9),(6),(215),(8),(219),(223),(225)*.

#### • Technique (anti-pathogen transgenesis)

A subsequent blockage or reduction of pathogen transmission was reported in *Aedes aegypti (208),(211),(212),* in *Anopheles gambiae (204),(213),(222), Anopheles stephensi (207),(214),(221),* and in the silkworm *Bombyx mori (225)*. However, in one publication the *Plasmodium falciparum* protection induced by an anti-malarial gene was inconsistent *(213)*.

#### • Fitness (anti-pathogen transgenesis)

In some cases, the anti-pathogen transgene led to no or low fitness cost in the modified insects *(208),(222),* thus allowing the modified insect to be reproductively competitive against their natural counterparts. However, anti-pathogen gene insertion also caused fitness benefits in *Anopheles gambiae* (*206)*, and *Anopheles stephensi* that fed *Plasmodium*-infected blood *(207),(221).* Since these outcomes increase vectors abundances and vectorial capacity, they constitute reversal outcomes.

#### • Release (anti-pathogen transgenesis)

Several publications, all of them computational modelling or laboratorial studies, estimated successful release of insects modified with an anti-pathogen transgene *(7),(207),(4),(220),(221),(224).* Some out oh those described the efficacy of different gene drives (which bias the inheritance of a particular gene to quickly and irreversibly spread it through a population) such as, Multi-locus assortment *(209)*, Medea and Killer-Rescue *(220*), and *Wolbachia (7),(4),(9),(5).* Two publications presented comparative studies describing advantages and disadvantages of several gene drives *(6),(8)*. Numerous computational modellings studies suggested a hard compromise between invasiveness and confinement (a high migration rate required to become established in neighboring populations, and low frequency persistence in neighboring populations for moderate migration rates) *(209),(220),(7),(4),(9),(5),(6),(8). Wolbachia* was referred as an efficient gene drive in some studies *(9),(7),(5),(4)* but according to Marshal and Hay, 2012 *(8*), was not reliable for confinement properties. Semele, Merea and two-locus engineered underdominance were the most promising in confinement properties and required lower introduction frequencies (compared to *Wolbachia*, Medea, single-allele underdominance, single-locus engineered underdominance and killer-rescue) *(8)*. Multi Locus Assessment, despite being less effective as gene drive, allows the test of ecological components before releases with more invasive gene drives *(209)*.

#### • Modification and Technique (lethal transgenesis)

Successful insertion of a lethal transgene was reported in *Aedes aegypti (226), Drosophila melanogaster (230)* and *Ceratitis capitata (228)* and subsequent lethality was laboratorial confirmed in *Aedes aegypti (226),* and *Drosophila melanogaster (230)*.

#### • Fitness (lethal transgenesis)

One publication reported no fitness costs caused by the modification on the modified insect and confirmed that the lethal transgene did not affect insecticide susceptibility *(228)*.

#### • Release (lethal transgenesis)

Computational modelling studies reported that elimination of vector insects might be an unrealistic objective. However, substantial suppression can nonetheless be achieved in certain conditions, such as an uniform spatial pattern and multiple lethal elements *(227)*, or a certain release ratio and population size *(229),(224)*. Elimination of a natural population after the release of insects with a dominant lethal was though reported in semi field studies *(228),(226)*.

#### • Long-term (lethal transgenesis)

One computational modelling study suggested lethal transgenesis long-term efficacy to be compromised by invasion of wild type insects *(224)*.

From all publications in this section, only one publication reported a field study, describing the ability to mate and copulate of a radiated insect, modified by a sterilizing technique (SIT) *(231)*. Also only three publications reported reversal outcomes, related to fitness benefits (as above mentioned). Main effective outcomes of transgenic and other non-symbiont-based modified insect vectors are presented in Table 3 (the complete data is presented in S2 Table).

**Table 3.**
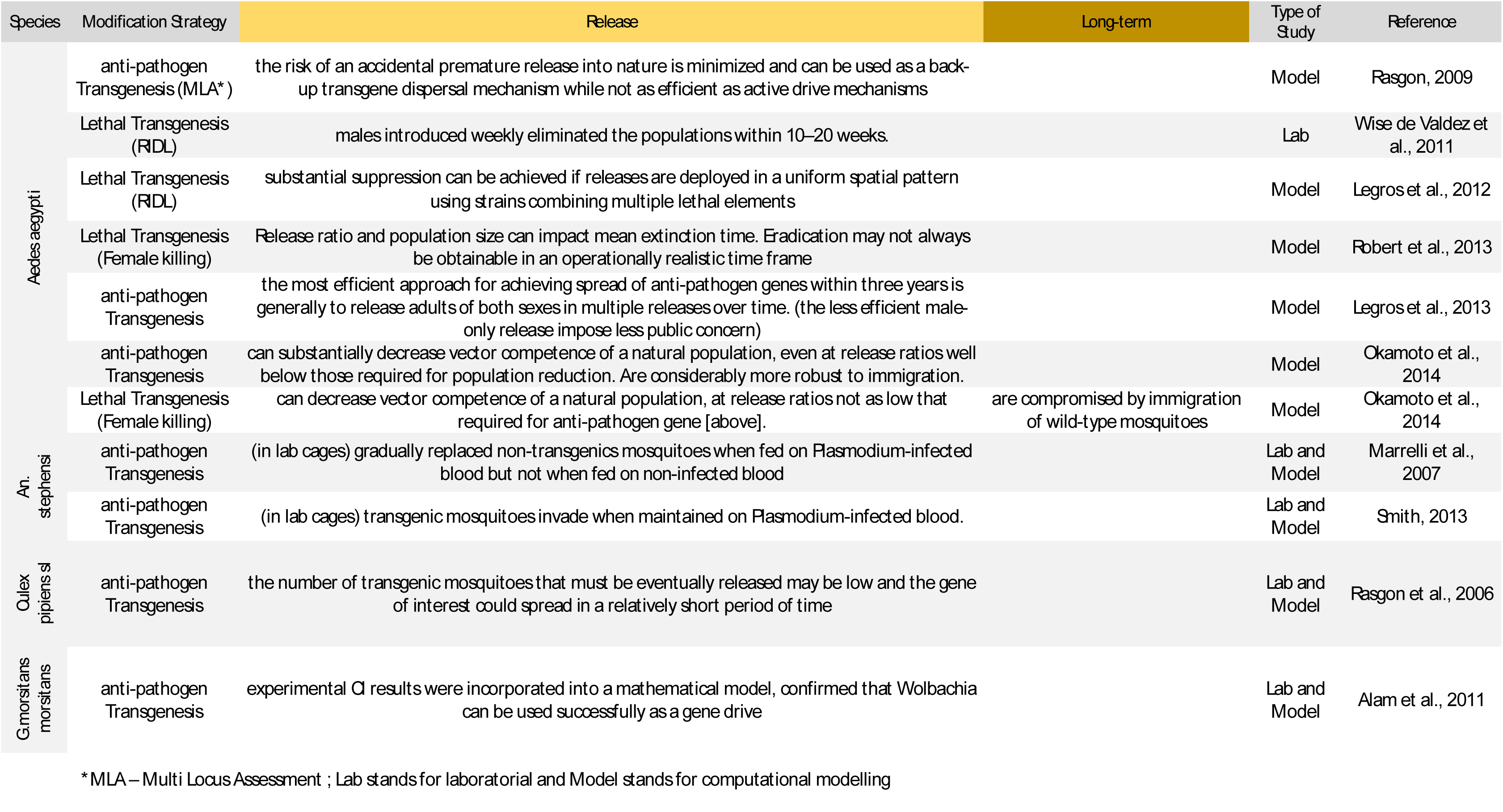
Main effective outcomes of transgenesis and other non-symbiont-based insect modification (vector insect species) Publications reporting effective outcomes on the topics: release, epidemiology and long-term (the ones closer to the strategy’s aim, that is transmission blockage or vector suppressing). Ineffective outcomes of the mentioned publications are also presented (in italic).

Five articles (2.1% out of the 237 with efficacy outcomes) contributed to the efficacy of insect modification as a vector control approach, regardless of the modification strategy used (see S8 Table). They reported results as diverse as: effective releases, in what concerns numbers of insects, and sex ratio, (*233*), the impact of laboratory rearing *(234)*, or descriptions of gene flow of eventual release sites *(235-237)*. Interestingly, in all analyzed publications regarding gene flow, no isolation was found between Islands and mainland, neither in Society Islands of French Polynesia *(236)*, nor Lake Victoria in Western Kenya *(235)*, nor in Bijagós archipelago in Guiné-Bissau (*237)*.

## EFFECTS

Apart from the intended effective traits, modifications also induced other effects into the modified insects, as reported in 155/44.4% publications (nTotal=349). Modifications induced effects at several levels: (i) at the specimen level, *i.e.* physiological effects such as, reproduction, immune response or microbiome of the modified insect; (ii) at the insect species level, concerning its evolution and/or behavior; and (iii) at the ecosystem level, affecting any other organism of the modified insect ecosystem (Fig 2 and Fig 7).

**Fig 7.**
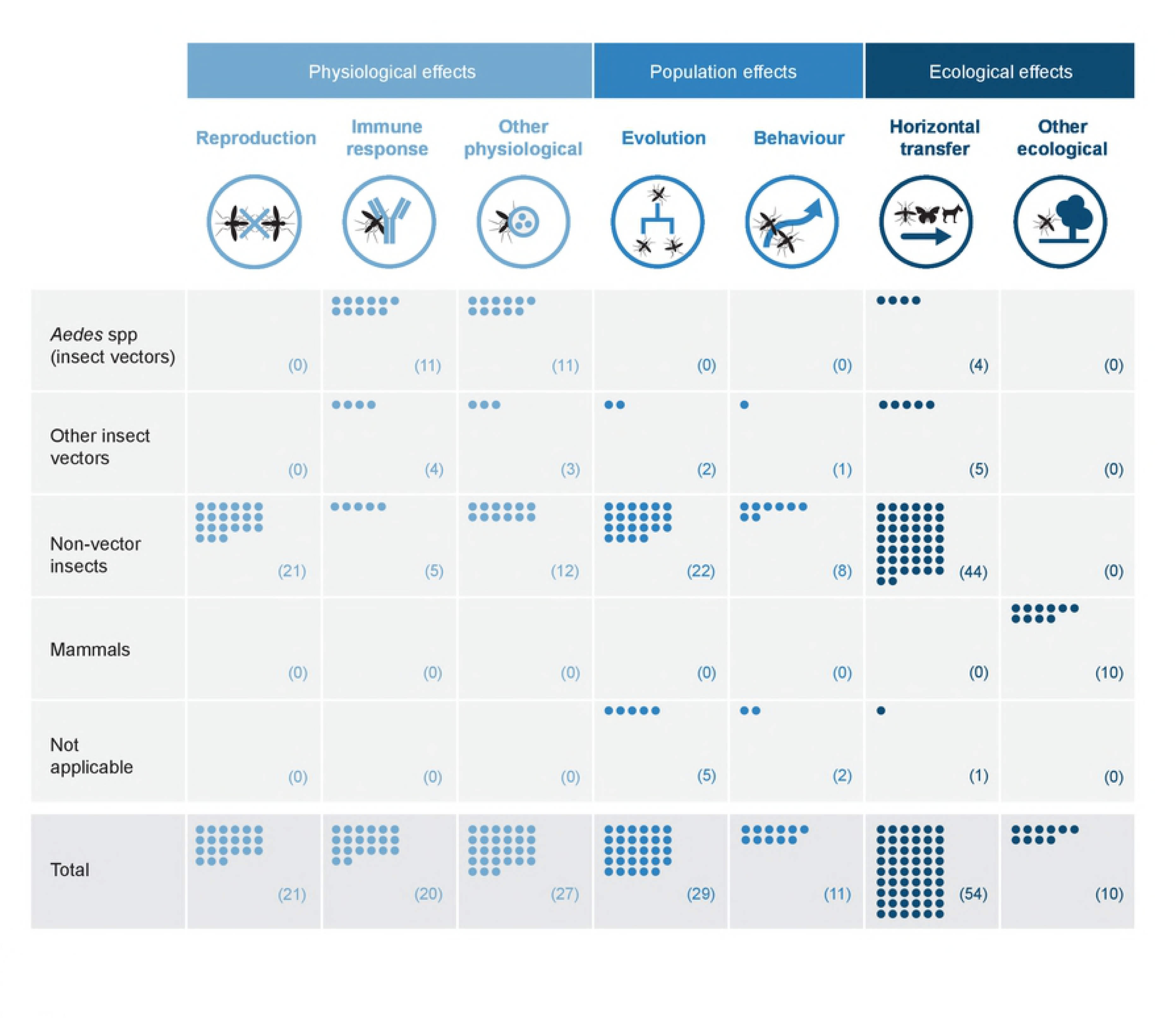
Number of publications reporting each Wolbachia-induced effect per taxonomic group

Majority of publications contributing to this theme covered effects specifically induced by *Wolbachia*. There were two exceptions only, all of them reporting populational effects: one publication describing effects on evolution, induced by other symbiont (*Rickestia*) *(1*), and one publication describing loss of assortative mating induced by laboratory rearing *(234)*.

### Physiological effects (at the insect specimen level)

A total of 63/18.1% publications (nTotal=349) reported that insect modification strategies may induce physiological effects on the target insect. Most frequent physiological effects found in analyzed publications were *Wolbachia*-induced effects on reproduction. The following reproductive modifications were reported: (i) male-killing (the death of male embryos during early embryonic development, with advantage for the surviving infected female siblings), stated in four non-vector species (*Ephestia Kueheniella –* butterfly, *Hypolimnas bolina-* butterfly, *Ostrinia scapulalis moth,* and *Tribolium madens - beetle*) *(53),(54),(155),(238-240);* (ii) feminization (the conversion of genetic males into functional females), described in two non-vector species (*Eurema hecabe*,and *Zyginidia pullula*) *(241-243),(172);* and (iii) parthenogenesis (the exclusive participation of females on reproduction, and production of female offspring), reported in several species of parasitoid wasps *(173),(244),(245).* Although cytoplasmic incompatibility (CI) can be considered a *Wolbachia* reproductive effect, since it is a required trait for *Wolbachia* use as control strategy, it was herein considered an efficacy trait rather than a reproductive effect (see CI results on Technique topic). Moreover, effects on reproduction also included sex ratio alterations *(246),(157),(247),(248),* exceptional sex mosaics *(168),* requirement of *Wolbachia* for oogenesis *(130),(131),* or changes in expression of genes associated with reproduction *(249),(172)*.

*Wolbachia* also induced effects on the immunity of the modified insect mainly through the up-regulation of effector genes. Report of *Wolbachia*-mediated induction of immune system was described in *Aedes aegypti (14),(90),(250),(187),(251),(23), Aedes albopictus (252),(253),(35), Aedes polyniensis (37), Anopheles gambiae (254),(253),(57), Anopheles stephensi* (*40*), and *Drosophila melanogaster (250),(249). Wolbachia* also induced reduction of the immune response by decreasing the ability to encapsulate parasitoid eggs in *Drosophila simulans (161) or* decreasing the ability to produce lead peroxides *(255)*.

Other physiological effects were also reported, such as, alteration in the insect’s microbiome, *(164),(138),(150),(39),(256),* gene expression (genes, microRNA, sRNA, or epigenetic effects), *(172),(257),(249),(258-260)*, or in its nutrition and metabolic mechanism *(261),(170),(71),(262-265),(129),(249),(266),(267),(137),(195),* see all information regarding physiological effects (also *(268),(269),(182),(270),(189),(271)*) in S4 Table.

### Populational effects (at insect population level)

According to 40/11.5% analyzed articles (nTotal=349), *Wolbachia* also affected its host population in several ways such as, altering its mitochondrial DNA (mtDNA) pattern, interfering in speciation process, on its behavior ecology or others. Changes in mtDNA patterns were reported in several non-vector insects (*272-283*), and in the vector mosquito *Culex pipiens (7).* Phylogenetic analysis of *Culex pipiens* complex populations from three continents indicated a *Wolbachia*-induced drastic reduction of mitochondrial variability, thus profoundly interfering in its population structure *(7)*. Several articles suggested that *Wolbachia* induced speciation, altering genomic diversity *(284)*, leading to premating behavioral isolation *(285),(286)* or to reproductive divergence (due to variable phenotypic effects) (*10*). *Wolbachia*-induced behavioral isolation is more likely in diploid and haploid than in haplodiploids hosts *(287)*, and can be more evident in hybrid zones (*288*). Examples of behavioral changes, all of them reported in non-vector insects or suggested by computational modelling studies, are the sex-role inversion on reproductive ritual *(246),* the increase of sexual promiscuity *(159),(289),* and the irreversible loss of sexual reproduction *(290),(173),(291)*.

Symbiont-induced speciation was also reported in *Neochrysocharis formosa* infected with *Rickestia* (*1)*. All data regarding populational effects (also *(292),(293),(160),(170),(247),(294), (172),(295),(144),(296-299))* is presented in S5 Table.

### Ecological effects (at insect ecosystem level)

Finally, 64/18.3% publications reported that *Wolbachia* is also able to induce alterations in other organism rather than its host, interfering thus, with host ecosystem. The majority of the publications covering this type of effect described the report or the estimation of horizontal transfer events (*i.e*. transfer between neighboring contemporary species) of genetic material, such as a gene or a symbiont. Horizontal transfers (HT) can occur through bacteriophages, parasitoids, hemolymph, etc. Reported HT events comprise innumerous type of transfers such as between symbionts *(300),(292),(301), (302-304),(137),* from *Wolbachia* to insect vector *(305-309)*, from *Wolbachia* to nematode species (*305)*, from insect to insect *(155),(116),(310-312),(118),(313),(294),(107),(314),(248),(315-320),(296),(321-324),(201),* and from insect to non-insect species *(325),(326)*, insect to bacteria (*327)*. All data regarding HT (also *(328),(329),(119),(167),(305),(330),(122),(331),(302),(245),(325),(332),(333),(307),(334-339)*) is described in S6 Table.

Apart from HT, *Wolbachia* can reach other organisms also inducing ecological effects. All publications reporting non-HT ecological effects consisted in studies in mammals. Those had contact with *Wolbachia* mainly *via* filarial infection. A *Wolbachia*-infected filarial nematode induced or exacerbates the filarial diseases pathogenesis such as, human subcutaneous dirofilariasis (*340)*, onchocerciasis (river blindness) *(341),(342)*, and lymphatic filariasis *(343),(344)* (Table 4).

**Table 4.**
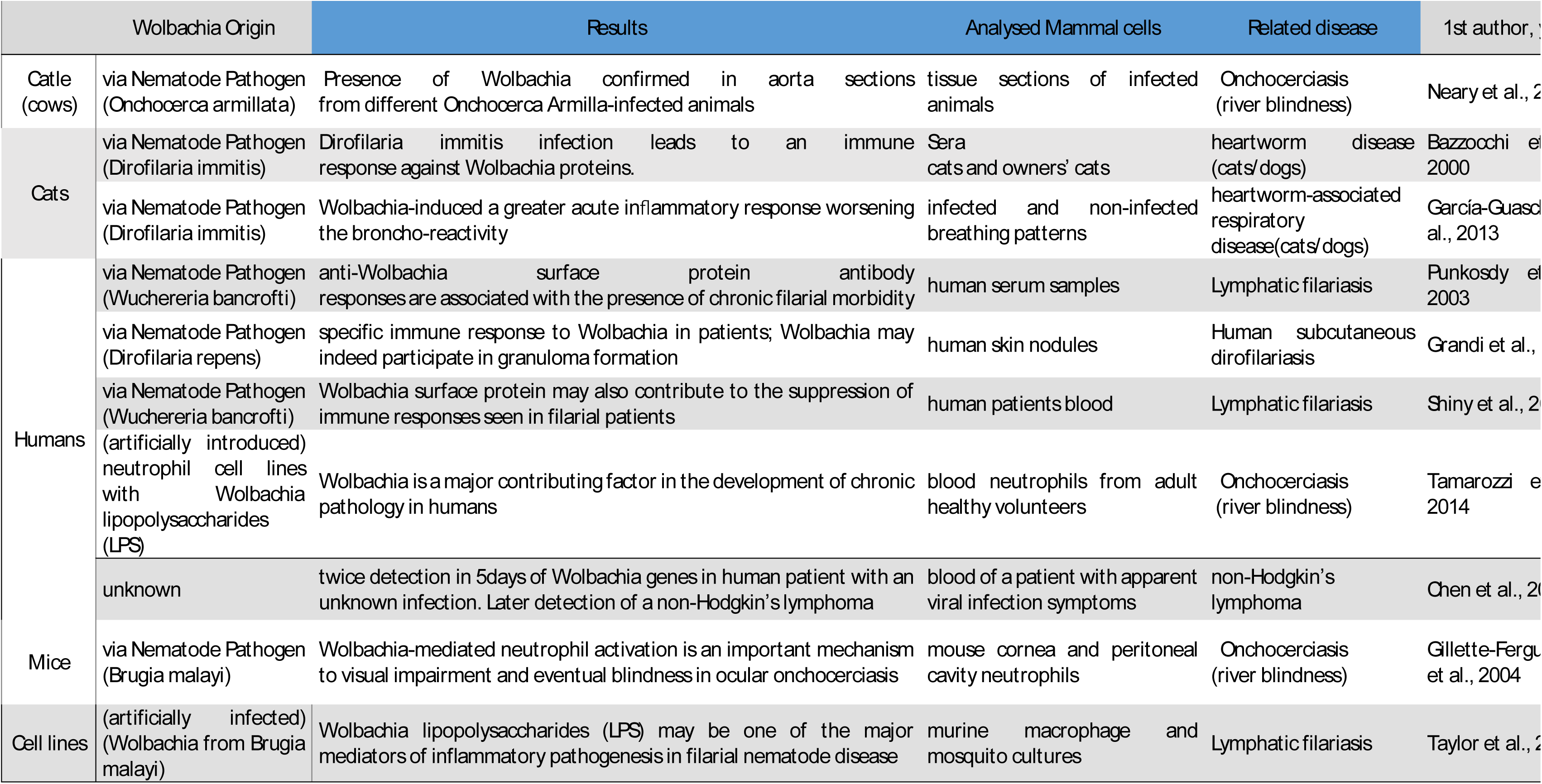
Other ecological effects: publications reporting Wolbachia-induced effects on mammals and respective main results

## DISCUSSION

To our knowledge, the present work constitutes a unique review on modified insects for vector-borne disease prevention, for several reasons. Rather than being the perspective of an author or a summary of the authors’ selection of publications, this review followed a structured methodological procedure. Furthermore, authors from present review are not involved in scientific projects to modify insects having, thus, no conflict of interest in the outcomes of this reflection. Moreover, present review enclosed several types of modifications in any vector or non-vector insect species, covering an uncommon comprehensiveness, and thus, offering an exceptional opportunity to observe trends and to outline the big picture of the modified insects. In that sense, this review constitutes a baseline of knowledge that not only can be fed with forthcoming publications to follow up trends, but also point out to questions that may need to be further explored. Additionally, this review developed a framework of themes, topics and outcomes, that organizes the extensive information available. Also, it goes beyond modifications’ efficacy, most commonly covered in current reviews (26)(23), exploring research articles also related to modifications’ effects and even the variability of the efficacy outcomes in different settings. Finally, it is to our knowledge the sole review on the subject gathering publications on mammals and exploring the eventual effects of insects’ modifications on this important taxonomic group where humans are included. According to the epidemiology definition, efficacy determines whether an intervention produces the expected result under ideal circumstances, while effectiveness measures the degree of beneficial effect under actual settings(38). This works explored efficacy (modification, technique and fitness) and effectiveness related topics (release, epidemiology, long-term and variability). However, to simplify classification, ‘efficacy’ was used as a broad term that included all above-mentioned topics.

Efficacy of the modifications was the most covered theme, being analyzed in 237 out of the 349 publications included in the review. Nevertheless, the amount of publications regarding modifications’ efficacy does not reflect the extent neither the robustness of the evidence available regarding it. Out of those 237, only 41 publications reported main effective outcomes: successful releases of modified insects, its positive epidemiological impact and/or its long-term efficacy (the remaining cover ineffective or primary effective outcomes). Out of the three main efficacy topics, the epidemiological impact was the less covered, reported in only five publications, none of them based on the most robust field studies, but all based on computational modelling studies instead. Main effective outcomes were obtained for several insect modifications, namely, *Wolbachia*-based strategies (replacement and suppression approaches), RIDL and female killing lethal transgenesis (both suppression approaches), computational modeled transgenesis regardless of the transgene (replacement and/or suppression approaches), and paratransgenesis (replacement approach). Seven publications (out of the 41) reported main effective outcomes based on semi-field or field studies, (corresponding to outcomes obtained with *Wolbachia* or RIDL strategies). Even though not many, these publications achieved critical outcomes: (i) field-released wMel-*aegypti* mosquitoes not only reached near fixation (despite a persistent low frequency of uninfected mosquitoes), but also maintained their effective traits such as CI, fixation and pathogen protection for at least two years after the release *(27)(28)* (from S1 Appendix) (ii) weekly introduction of *Aedes albopictus* males with a dominant lethal (RIDL) led to the eradication of a laboratory cages population in 10-20 weeks *(228)* (from S1 Appendix).

Present review also described reversal outcomes obtained after the release of transgenic insects or insects modified with *Wolbachia* (reported in 39 publications). Out of those, only one publication correspond to a field study, but in this case the release of modified insects may lead to an increase in the vector insect population particularly if occurring when its natural abundance is at its maximum *(100)* (from S1 Appendix).

Furthermore, analyzed publications also reported biological effects (at physiological, populational and ecological level) of *Wolbachia*-based insect modifications. Effects on mammals should be particularly and carefully explored. Monitoring of an eventual increase of lymphatic filariasis severity, non-Hodgkin’s lymphoma incidence, or unrecognized infections may be advised in areas already subjected to *Wolbachia*-insects releases.

Even though there were not found publications reporting eventual effects of transgenesis and other non-symbiont modifications, some of their effects were already discussed in the literature. In what concerns suppressing strategies, such as RIDL (lethal transgenesis), female killing (lethal transgenesis), SIT or RNAi-mediated sterilization, the elimination of a species leads to profound changes in its ecosystem, such as eventually putting some non-target species in risk or giving opportunity to not-targeting species to expand (35)(23). Moreover, some authors have been arguing that biodiversity loss may even be associated to emergence of vector-borne diseases (36) (37). Regarding replacement approaches, such as anti-pathogen transgenesis, the effects of the transgene in the ecosystem are unknown. When associated with *Wolbachia* as gene drive, there are emerging questions regarding the lateral transfer of the inserted transgene to *Wolbachia* itself or via *Wolbachia* to other organisms.

Since the inclusion criteria were restricted to studies on insects and mammals, results regarding ecological effects may be limited to these taxonomic groups. Horizontal transfer events of *Wolbachia* genetic material to non-insect non-mammal species were even though reported *(305),(325),(326*) (from S1 Appendix). It is not surprising that publications covering *Wolbachia* were the most founded in this review. Since *Wolbachia* is a natural bacterium, *Wolbachia*-based strategies are much more accessible for study than other patented strategies such as, RIDL or transgenesis. Moreover, besides those studies specifically oriented for a *Wolbachia*-based strategy, other studies regarding *Wolbachia* were included (mainly regarding natural *Wolbachia*) whenever their results contributed to the efficacy or the effects of *Wolbachia* as a vector control strategy. Furthermore, *Wolbachia* is a unique term while for other strategies several diverse terms may be used (such as, the name of the particular transgene), and may occur that not all were covered within the research expressions.

In what concerns the year of the publications it is clear how recent this topic is, being almost the totality been published in the last 20 years. This can, at least partially, explain why in several topics we found a gap of knowledge such as the long-term efficacy or the epidemiological impact of the modifications.

In conclusion, insects modifications strategies appear as a promising innovative alternative to overcome an unprecedented increase of vector-borne, mainly arboviral, diseases. Nevertheless, these modification tools still lack evidence on field-based efficacy mainly regarding epidemiological and long-term impact. Field releases in endemic areas could provide that kind of missing evidence. However, relevant questions remain without a solid understanding. Eventual reversal outcomes on disease transmission, or irreversible biological effects (including effects on mammals) need to be explored, dispelled or resolved. This leads for demand of studies before testing their definite effectiveness in the field. However, some of these questions could only have a robust answer if these strategies would be implemented, needing to take the risk to observe reversal outcomes and/or irreversible effects, in order to confirm the efficacy of these strategies. This reflect the current dilemma that is under the use of modified insects to prevent vector-borne diseases. The level of variability of existing evidence suggests the need to generate local/specific evidence in each setting of an eventual release. Importantly, available preventive strategies should not remain on hold while modified insects do not offer an effective and safe solution. A comprehensive cost-effectiveness analysis could be an important tool when deciding to proceed or not with these innovative strategies, and/or to improve other available strategies. Therefore, an adequate decision could be made for each particular setting, evaluating the pros and cons of these approaches and of each modifying technique.

## LIST OF SUPPLEMENTARY INFORMATION

S1 Checklist – PRISMA checklist

S1 Appendix – List of analyzed publications and summary of analysis

S2 Appendix - Research expressions, review assumptions and list of analyzed categories

S1 Fig - Quantitative description of the included publications according to: type of insect modification strategy (A), type of study (B), study’s species (C), year of publication (D)

S1 Table - Results regarding effective and ineffective outcomes of wolbachia and other symbiont-based insect modification strategy: a) on Insects vectors; b) on Non-vector Insects (from pp.24)

S2 Table - Results regarding effective and ineffective outcomes of Transgenesis and other non-symbiont-based insect modification strategy: a) on Insects vectors; b) on Non-vector Insects (from pp.7)

S3 Table-Results regarding effective and ineffective outcomes regardless of the insect modification strategy

S4 Table - Results regarding modification-induced physiological effects: a) on insect reproduction; b) on insect immune response (pp.3); c) other modification-induced physiological effects (pp.5)

S5 Table-Results regarding modification-induced populational effects (evolution and behavior) S6 Table-Results reporting horizontal transfer events in modified insects

## ACKNOWLEDGMENTS

We also would like to thank Carlos Rodrigues, Gil Costa, Gonçalo Seixas, Luis Xavier and Patrícia Salgueiro for their contributions.

